# Host-directed therapy with 2-Deoxy-D-glucose inhibits human rhinoviruses, endemic coronaviruses, and SARS-CoV-2

**DOI:** 10.1101/2022.05.24.493068

**Authors:** Laxmikant Wali, Michael Karbiener, Scharon Chou, Vitalii Kovtunyk, Adam Adonyi, Irene Gösler, Ximena Contreras, Delyana Stoeva, Dieter Blaas, Johannes Stöckl, Thomas R. Kreil, Guido A. Gualdoni, Anna-Dorothea Gorki

## Abstract

Rhinoviruses (RVs) and coronaviruses (CoVs) upregulate host cell metabolic pathways such as glycolysis to meet their bioenergetic demands for rapid multiplication. Using the glycolysis inhibitor 2-deoxy-D-glucose (2-DG), we assessed the dose-dependent inhibition of viral replication of minor- and major-receptor group RVs in epithelial cells. 2-DG disrupted RV infection cycle by inhibiting template negative-strand as well as genomic positive-strand RNA synthesis, resulting in less progeny virus and RV-mediated cell death. Assessment of 2-DG’s intracellular kinetics revealed that after a short-exposure to 2-DG, the active intermediate, 2-DG6P, is stored intracellularly for several hours. Finally, we confirmed the antiviral effect of 2-DG on pandemic SARS-CoV-2 and showed for the first time that 2-DG also reduces replication of endemic human coronaviruses (HCoVs). These results provide further evidence that 2-DG could be utilized as a broad-spectrum antiviral.

**HIGHLIGHTS:**

- 2-DG inhibits replication of minor- and major-group rhinoviruses in epithelial cells including human nasal epithelial cell.
- 2-DG disrupts rhinovirus infection cycle and reduces rhinovirus-mediated cell death *in vitro*.
- 2-DG treatment attenuates viral load of endemic coronaviruses *in vitro*.

## INTRODUCTION

Rhinoviruses (RVs) and endemic human coronaviruses (HCoVs) are the major cause of acute respiratory tract (RT) infections in humans [1], [2]. These are largely self-limiting in healthy adults, where they usually remain confined to the upper respiratory tract. However, as the viruses spread rapidly and circulate seasonally, they lead to high incidence rates on an annual basis. These can cause severe morbidity in elderly, children, and immune-compromised patients [3]–[6]. Along with human suffering, these viral infections lead to high economic losses and healthcare costs [7], [8]. While global efforts are underway to develop an effective therapy, the current lack of FDA-approved antivirals has limited the treatment of RT infections to supportive and symptomatic care.

As *Picornaviridae*, RVs are non-enveloped and contain a positive-sense single-stranded RNA genome ((+)ssRNA) [9]. They are divided into three species, RV-A, RV-B and RV-C. RV-A and RV-B are further classified as minor- and major-group based on the cognate host cell receptors they use for cell entry [10]–[12]. Coronaviruses (CoVs) are enveloped viruses, belong to the *Coronaviridae* family and also contain a (+)ssRNA genome [13]. They are classified into four major genera: alpha, beta, gamma, and delta, targeting a variety of host species. In humans, strains from the alpha [14]–[16] and beta genera [17] are known to induce common colds similar to the ones caused by RVs [18], [19]. However, three strains from the beta genus, including Severe Acute Respiratory Syndrome Coronavirus 2 (SARS-CoV-2) were found to be more pathogenic with high fatality rates [20].

Viruses are dependent on the host cell metabolism and host cell machinery to ensure their replication. RVs and CoVs in particular are known to hijack and reprogram the host cell metabolic pathways for rapid multiplication, causing an increase in bioenergetic demand [21], [22]. This leads to an elevated anabolic state, forcing the host cell to synthesize more lipids and nucleotides using glucose and glutamine as substrates [23]. In addition, there is an increased demand for energy in the form of adenosine triphosphate (ATP) for viral replication and assembly, which is predominantly provided by glycolysis [23]–[25]. As an essential metabolic pathway, this involves breakdown of hexoses like glucose into pyruvate for ATP production. This dependency of RVs and CoVs, and presumably other viruses on host glucose metabolism for replication presents a promising target for the development of effective antiviral therapies.

2-Deoxy-D-glucose (2-DG), a stable analogue of glucose, is taken up by cells via glucose transporters and subsequently phosphorylated to 2-deoxy-D-glucose-6-phosphate (2-DG6P) by hexokinase [26], [27]. Unlike in glucose metabolism, 2-DG6P cannot be further metabolized by phosphoglucose isomerase [28]. This leads to intracellular accumulation of 2-DG6P and arrest of glycolysis at the initial stage, causing depletion of glucose derivatives and substrates crucial for viral replication [29]. Previously, it has been demonstrated that 2-DG affects viral replication by reverting virus-induced metabolic reprogramming of host cells [24], [25], [30], [31].

The present study explores the broad-spectrum antiviral activity of 2-DG. In this process, we investigated the antiviral activity of 2-DG against minor- and major-group RVs in epithelial cells including primary human nasal epithelial cells (HNECs), the main site of RV replication. In concurrent experiments, we characterized 2-DG’s intracellular kinetics. Finally, to better understand the inhibitory activity of 2-DG on the RV infection cycle, we quantified the template (−)ssRNA as well as the genomic (+)ssRNA and analyzed 2-DG’s effect on RV-mediated cell death. Finally, we assessed the antiviral activity of 2-DG against endemic HCoVs as well as the pandemic SARS-CoV-2 strain. In summary, our study provides further evidence that reverting virus-induced metabolic reprogramming by 2-DG treatment critically affects viral RNA replication and thus holds great potential in combating respiratory viral infections.

## METHODS

Details including supplier and catalogue number of all materials used are listed in Supplement table 1.

### Cell culture

Cells were seeded in either 24-well tissue culture plates or T25 flasks and incubated at 37 °C in media and densities (cells per well or cells per flask) for the given times as indicated below; human nasal epithelial cells (HNECs) in HNEC medium (Pneumacult-ex plus basal medium supplemented with 1x Pneumacult-ex plus supplement, 0.1 % Hydrocortisone stock solution and 1 % Penicillin/Streptomycin (100 Units/mL) at 4.5×10^4^ cells/well (72 h) and HeLa Ohio cells in HeLa Ohio medium (RPMI 1640 medium supplemented with 10 % fetal bovine serum (FBS), 1 % Penicillin/Streptomycin (100 Units/mL) and 2 mM L-glutamine) at 2×10^5^ cells/well (16-20 h). LLC-MK2 and MRC-5 cells were cultured in T25 cell culture flasks in LLC-MK2 medium (Eagle-MEM supplemented with 10 % FBS, 1x non-essential amino acids solution (NEAS), 100 mg/mL Gentamycin sulfate and 25 mM HEPES) and MRC-5 medium (Eagle-MEM supplemented with 10 % FBS, 2 mM L-Glutamine, 1x NEAS, 1 mM sodium pyruvate, 100 mg/mL gentamycin sulfate, 0.15 % sodium bicarbonate) at densities of 8×10^5^ and 9×10^5^ cells/flask, respectively. Vero cells were cultivated in TC Vero medium (supplemented with 5 % FBS, 2 mM L-Glutamine, 1x NEAS, 100 mg/mL gentamycin sulfate, and 0.075 % sodium bicarbonate).

### Viral infection and 2-DG treatment

HeLa Ohio cells and HNECs were infected for 1 h at 37 °C or 34 °C with RV at 0.005 to 0.5 TCID_50_/cell and 4.5×10^4^ TCID_50_/well, followed by treatment with 2-DG for 6 h, 24 h or 48 h. The supernatant from the cells was then subjected to virus titer analysis or the cells were treated with cell lysis buffer for RNA extraction. LLC-MK2 cells and MRC-5 cells were infected with SARS-CoV-2 (Beta-CoV/Germany/BavPat1/2020) (MOI of 0.001) at 36 °C and HCoV-229E (MOI of 0.01) at 36 °C or HCoV-NL63 (MOI of 0.01) at 33 °C, respectively. Cells were treated with 2-DG 1 h post-infection and samples were collected at the indicated times for virus titer analysis.

### RNA isolation and cDNA synthesis

Intra- and extra-cellular RNA was isolated according to the ExtractMe Total RNA Kit instructions. To avoid bias in extracellular RNA isolation, an internal spike-in RNA control was added to each sample. RNA concentration and purity was assessed using a nanophotometer. cDNA was synthesized according to the First strand cDNA synthesis kit using the program: 37 °C for 60 min and 70 °C for 5 min. Measurement of viral negative-sense single-strand RNA ((−)ssRNA) was performed as previously described [32] except that the synthesized cDNA wasn’t RNase treated and purified. The cDNA from (−)ssRNA was synthesized using a mix of strand-specific, chimeric sequence-containing primer chimHRV-b14_RT and control primer HPRT_R (Supplement table 1) instead of oligo(dT).

### qPCR

qPCR was performed using SYBR green mix and primers as specified in Supplement table 1. For measuring intracellular viral RNA, gene expression was normalized to HPRT using the Livak method [33] and expressed as fold change to control (infected, but untreated). Primers HRV-B14_R and chimHRV-b14_R1 were used for measurement of viral (−)ssRNA. For extracellular viral RNA, synthetic oligo standard (HRV-B14_F, HRV-B14_R and HRV-B14 primer amplicon, Supplement table 1) was used to generate a standard curve for the calculation of viral copy number by interpolation. Based on the qPCR data, the IC_50_ was calculated using least square regression on Prism 9.0.2.

### Virus titration

Samples from SARS-CoV-2, HCoV-229E and HCoV-NL63 were titrated on Vero cells, MRC-5 cells, and LLC-MK2 cells, respectively. Samples from RV-B14 were titrated on HeLa Ohio cells. Titration was performed using eightfold replicates of serial half-log_10_ (for SARS-CoV-2, HCoV-229E and HCoV-NL63) or log_10_ (for RV-B14) dilutions of virus-containing samples followed by incubation at 36 °C (SARS-CoV-2, HCoV-229E), 33 °C (HCoV-NL63) and 34 °C (RV-B14) for 5-7 days (SARS-CoV-2, HCoV-229E, RV-B14) or 9-11 days (HCoV-NL63). Wells were inspected under a microscope for cytopathic effect (CPE). For RV-B14, CPE was visualized by crystal violet staining. Recognizable CPE at each tested dilution was used to determine the dose according to Reed and Muench [34] and reported as log_10_-transformed median tissue culture infectious dose per milliliter (log_10_[TCID_50_/mL]).

### Virus-induced cytopathic effect

HeLa Ohio cells were infected for 1 h at 37 °C with RV-B14 (0.5 TCID_50_/cell) followed by 2-DG treatment for 24 h or 48 h at 37 °C. CPE was visualized by crystal violet staining. The effect of 2-DG on virus-induced cell death was assessed by calculating the ratio of the average of treated, uninfected to each treated, infected sample value.

### Cell viability

HNECs were treated with 2-DG for 7 h at 37 °C. Cell viability was assessed by crystal violet staining. The effect of 2-DG on cell viability was calculated relative to untreated cells.

### Crystal violet staining

Cells were incubated with crystal violet solution (0.05 % crystal violet in 20 % methanol) for 30-60 min, washed with ddH_2_O, air-dried, followed by 25 % glacial acetic acid. The absorbance was recorded at 450 nm.

### Glucose-uptake assay

Cells were treated with 2-DG in the absence of glucose for 10 min at 37 °C, followed by washing with PBS and incubation for up to 270 min in glucose-free medium. 2-DG uptake was assessed using the Glucose-Uptake Glo™ Assay kit. Luminescence was recorded on a microplate reader. 2-DG6P levels were calculated as percentage of signal upon exposure to 2-DG after subtracting the background value obtained from control samples (not treated with 2-DG).

### Statistical analysis

The graphs show pooled results of independent experiments with each experiment containing two to four cell culture wells per condition with the standard error of the mean (SEM). Analysis of statistical significance was performed using Student’s *t*-test (unpaired analysis) or ordinary one-way ANOVA with Dunnett’s correction or 2-way ANOVA with Bonferroni’s correction and considered significant when p < 0.05 (*p ≤ 0.05, **p ≤ 0.01, ***p ≤ 0.001, ****p ≤ 0.0001).

## RESULTS

### 2-DG inhibits RV replication in HeLa Ohio cells and HNECs

2-Deoxyglucose (2-DG) treatment has been shown to inhibit rhinovirus (RV) infection by reverting RV-induced anabolic reprogramming of host cell metabolism [25]. While the effect of 2-DG on RV-B14 [25] and RV-C15 [35] was shown before, its effect on additional serotypes belonging to minor- and major-group RVs [10]–[12] remains to be investigated. For this, HeLa Ohio cells were infected with minor-group (RV-A1B, RV-A2) and major-group (RV-A89, RV-A16, RV-A54) RVs. 2-DG treatment led to a dose-dependent reduction in intracellular viral RNA levels of all major- and minor-group RVs tested (Supplement Figure 1). As 2-DG is transported into cells utilizing the same transporters as glucose, this results in a competition for the uptake of 2-DG [26], [27]. The glucose concentration in conventional cell culture media ranges from 2 g/L to 4.5 g/L and is much higher compared to *in vivo* glucose levels (e.g., in the blood it ranges from 3.9 to 5.6 mmol/L i.e., 0.7 to 1 g/L). Therefore, we tested the antiviral effect of 2-DG under physiological glucose levels (Figure 1). We reduced the glucose concentration in the cell culture medium to 1 g/L to mimic a setting corresponding to human plasma. 2-DG treatment at physiological glucose levels showed an even stronger inhibitory effect on intracellular viral RNA levels of all major- and minor-group RVs (Figure 1A). With the highest tested concentration of 2-DG (30 mM) we observed a complete abolishment of viral RNA replication (Figure 1A). In line with these results, the absolute half-maximal inhibitory concentration (IC_50_) of 2-DG was lower under the physiological glucose setting: The IC_50_ ranged from 1.92 mM to 2.67 mM as compared to 3.44 mM to 9.22 mM for cells infected and treated under conventional culture conditions (Figure 1B, Supplement table 2).

**Figure 1:**
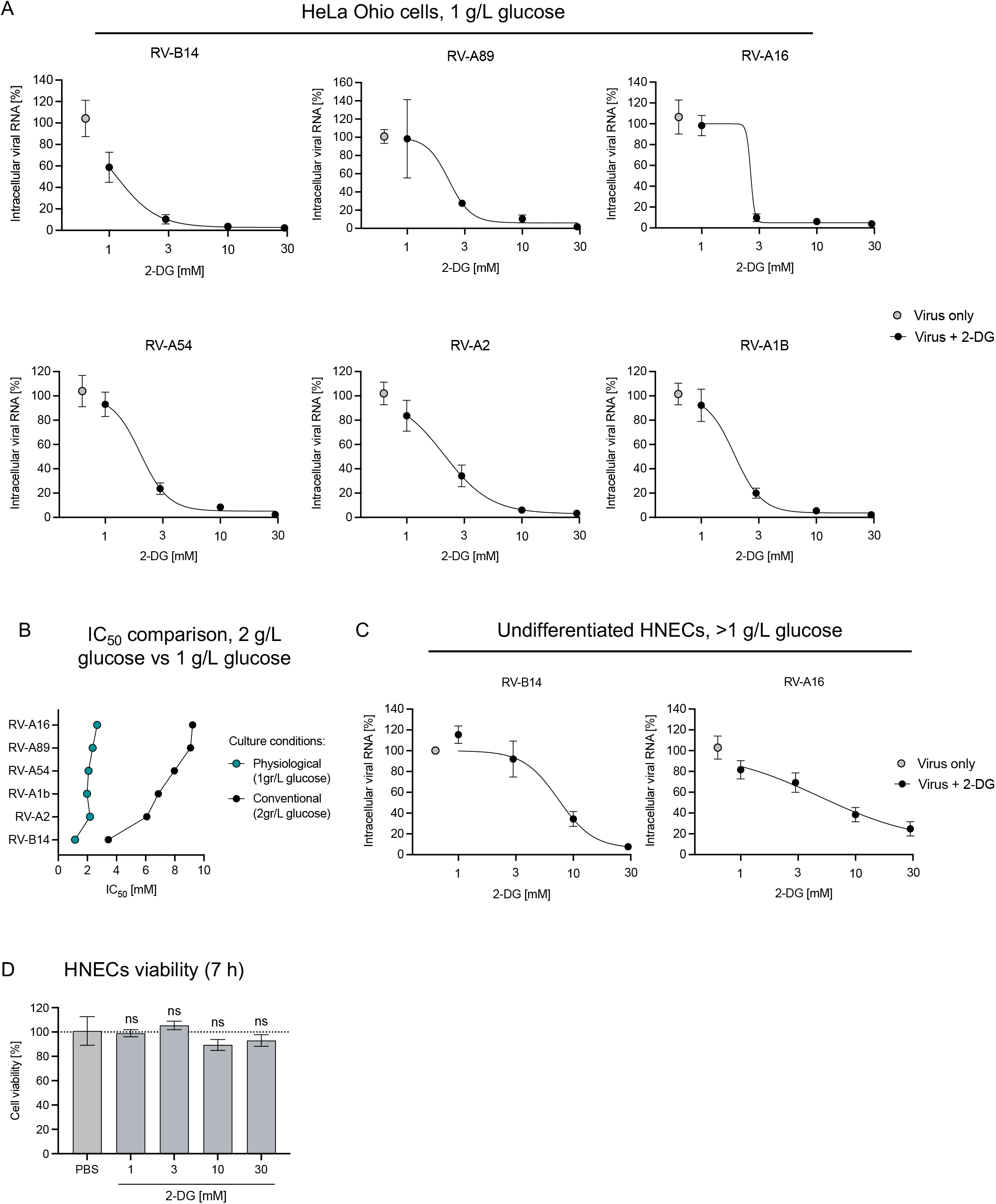
Inhibition of RV replication by 2-DG in HeLa Ohio cells and HNECs. Intracellular viral RNA was assessed by qPCR 7 h post-infection at 0.005 TCID_50_/cell for the indicated RV strains in HeLa Ohio cells in medium containing 1g/L glucose (A). Comparison of IC_50_ of 2-DG on the indicated RV strains under physiological versus conventional culture conditions (B). Intracellular viral RNA was assessed by qPCR 7 h post-infection at 4.5×10^4^ TCID_50_/well for the indicated RV strains in HNECs (C). In (A) and (C) cells were treated with the indicated concentrations of 2-DG (represented on a log10 scale) 1 h post-infection until samples were collected. The viability of HNECs was assessed at 7 h post-treatment with indicated concentrations of 2-DG (D). Graphs show pooled result ± SEM of 3-4 independent experiments. HNEC: human nasal epithelial cells, RV: rhinovirus.

Further, we evaluated the effect of 2-DG on RV-B14 and RV-A16 replication in human nasal epithelial cells (HNECs), the natural replication site for RVs. In line with the previous findings, 10 mM and 30 mM 2-DG treatment strongly inhibited RV-B14 and RV-A16 replication (Figure 1C). To be noted, unlike in HeLa Ohio cell culture medium, where the glucose level is known, glucose levels in HNECs culture medium (STEMCELL Technologies) are not disclosed.

As 2-DG inhibits glycolysis, a major energy generating pathway, we assessed whether it has an impact on cell viability in our setting. We did not measure a significant reduction in cell viability after 7 h 2-DG treatment (Figure 1D). Taken together, the data suggests that 2-DG inhibits RV replication in a dose-dependent manner, independent of the RV strain and cell type used. No toxic effects on the cells were recorded at concentrations those employed in the virus inhibition experiments. Furthermore, we observed better uptake and enhanced antiviral activity of 2-DG at physiological glucose levels.

### A short exposure to 2-DG leads to extended intracellular storage of 2-DG6P

Once 2-DG is taken up by the cell, it is phosphorylated to 2-deoxy-D-glucose-6-phosphate (2-DG6P), which leads to the arrest of glycolysis and altering of viral replication [25]. Thus, the kinetics of cellular uptake and intracellular storage are crucial for the antiviral activity of 2-DG. Therefore, we investigated the intracellular concentration kinetics of 2-DG6P in HeLa Ohio cells and HNECs. The experimental setup was designed to mimic treatment setting of 2-DG *in vivo*, e.g., a local application to the nasal cavity. Cells were treated with 1 mM and 10 mM 2-DG for 10 min, followed by washing to remove extracellular 2-DG and subsequent incubation up to 270 min and quantification of 2-DG6P levels (Figure 2A). At time zero (immediately after the 10 min 2-DG treatment), higher 2-DG6P levels were observed in 10 mM 2-DG treatment compared to 1 mM 2-DG treatment, in both HeLa Ohio cells and HNECs (Figure 2B, 2C, left graph). The intracellular 2-DG6P level measured at time zero was then set to 100 %, and the percentage decay of 2-DG6P over time was calculated. In HeLa Ohio cells 3.5 % ± 0.6 % (mean±SEM) and 18.5 % ± 3.4 % 2-DG6P were measured in 1 mM and 10 mM 2-DG treated cells after 270 min (Figure 2B). In the case of HNECs, higher levels of 2-DG6P retention were observed after 270 min; 10.1 % ± 1.5 % and 42.6 % ± 7.2 % 2-DG6P being detected in cells pre-treated with 1 mM and 10 mM 2-DG (Figure 2C), respectively. Collectively, the data suggest that short exposure of the cells to 2-DG leads to an intracellular accumulation of the active intermediate 2-DG6P for several hours.

**Figure 2:**
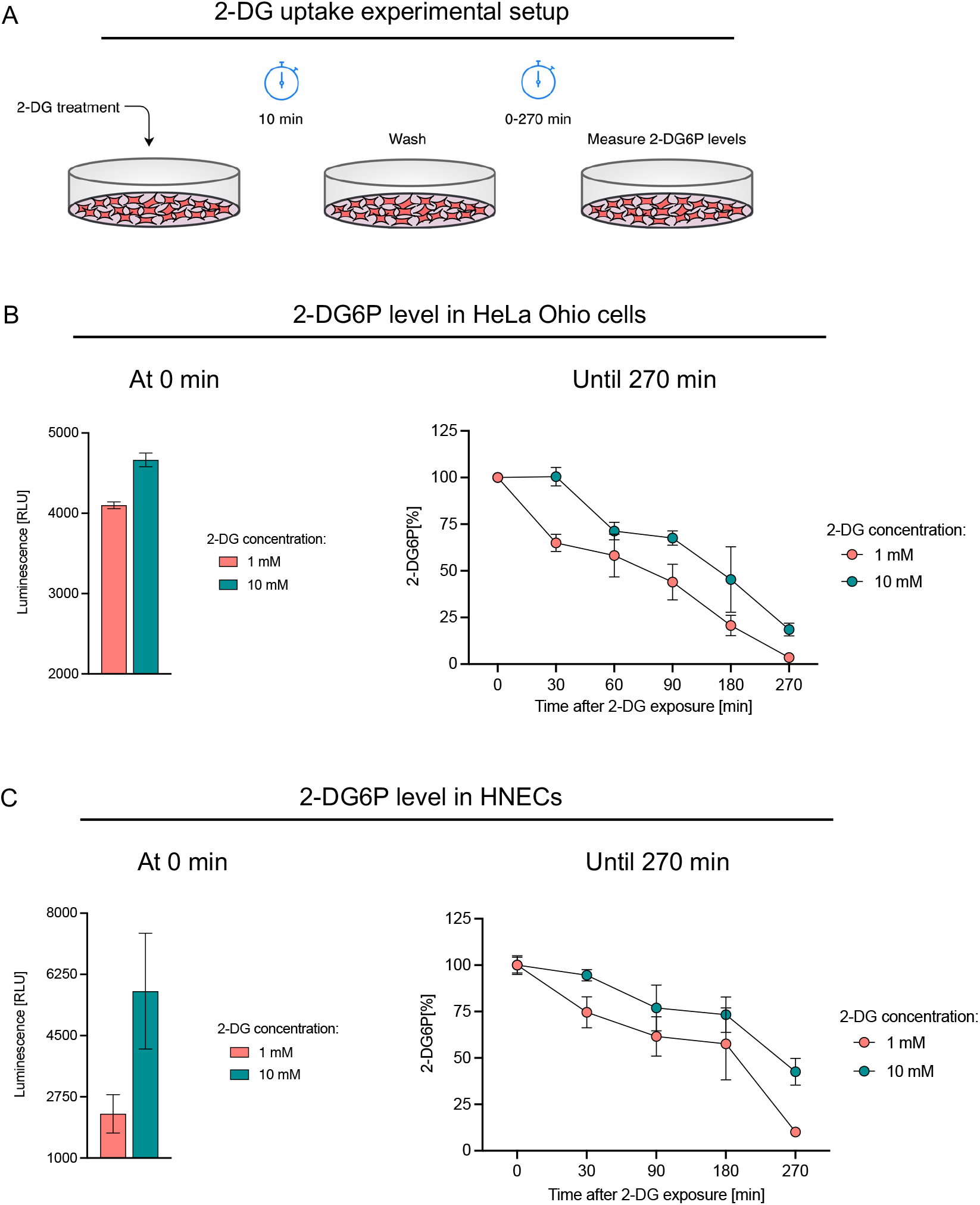
Intracellular storage of 2-DG6P after short-term exposure to 2-DG. 2-DG uptake experimental setup (A). Luminescence-based measurements of intracellular 2-DG6P at the indicated times after HeLa Ohio cells (B) or HNECs (C) were exposed to 2-DG for 10 min. In (B) and (C), the left graphs show the 2-DG6P levels (in RLU) at time 0 min (i.e., immediately after 10 min 2-DG treatment), and the right graphs show percentage decay of 2-DG6P over time in HeLa Ohio and HNECs, respectively. Data show pooled result ± SEM of 2-3 independent experiments. RLU: relative luminescence units, HNEC: human nasal epithelial cells.

### 2-DG disrupts RNA template strand synthesis and inhibits RV-mediated cell death

In our initial investigation of 2-DG mediated inhibition of RV replication, we measured the (+)ssRNA copies because of its abundance (10,000-fold higher than (−)ssRNA) [36] and the ease of quantification. However, the RV replication cycle involves generation of (−)ssRNA which is used as template for the replication of positive strand genomes [37]. Thus, the determination of (−)ssRNA serves as a means to quantify double stranded RNA (dsRNA), which is an intermediate of viral replication [36], [38]. Therefore, we analyzed the influence of 2-DG on synthesis of (−)ssRNA and of (+)ssRNA at 24 h post-infection. 10 mM 2-DG treatment led to a significant decrease in template (−)ssRNA levels of RV-B14 at 24 h post-infection (Figure 3A). This result was closely mirrored by decrease in the (+)ssRNA strand upon 2-DG treatment (Figure 3A). Simultaneously, we found that 2-DG treatment led to a significant decrease in the number of viral RNA copies in the supernatant (Figure 3B), implying an impairment of the amount of released virus. Next, we assessed 2-DG’s impact on viral load by means of median tissue culture infectious dose (TCID_50_) assays. RV-B14 infected HeLa Ohio cells were treated with 2-DG at 3.57 mM, corresponding to IC_90_, up to 48 h and the supernatants containing progeny virus were collected every 24 h and analyzed by virus infectivity assay. The above IC_90_ concentration of 2-DG was calculated from the previously derived dose-response curve in HeLa Ohio cells (Figure 1A, RV-B14). In comparison to the untreated cells, 2-DG treated cells showed a significant reduction in viral load 48 h post-infection (Figure 3C).

**Figure 3:**
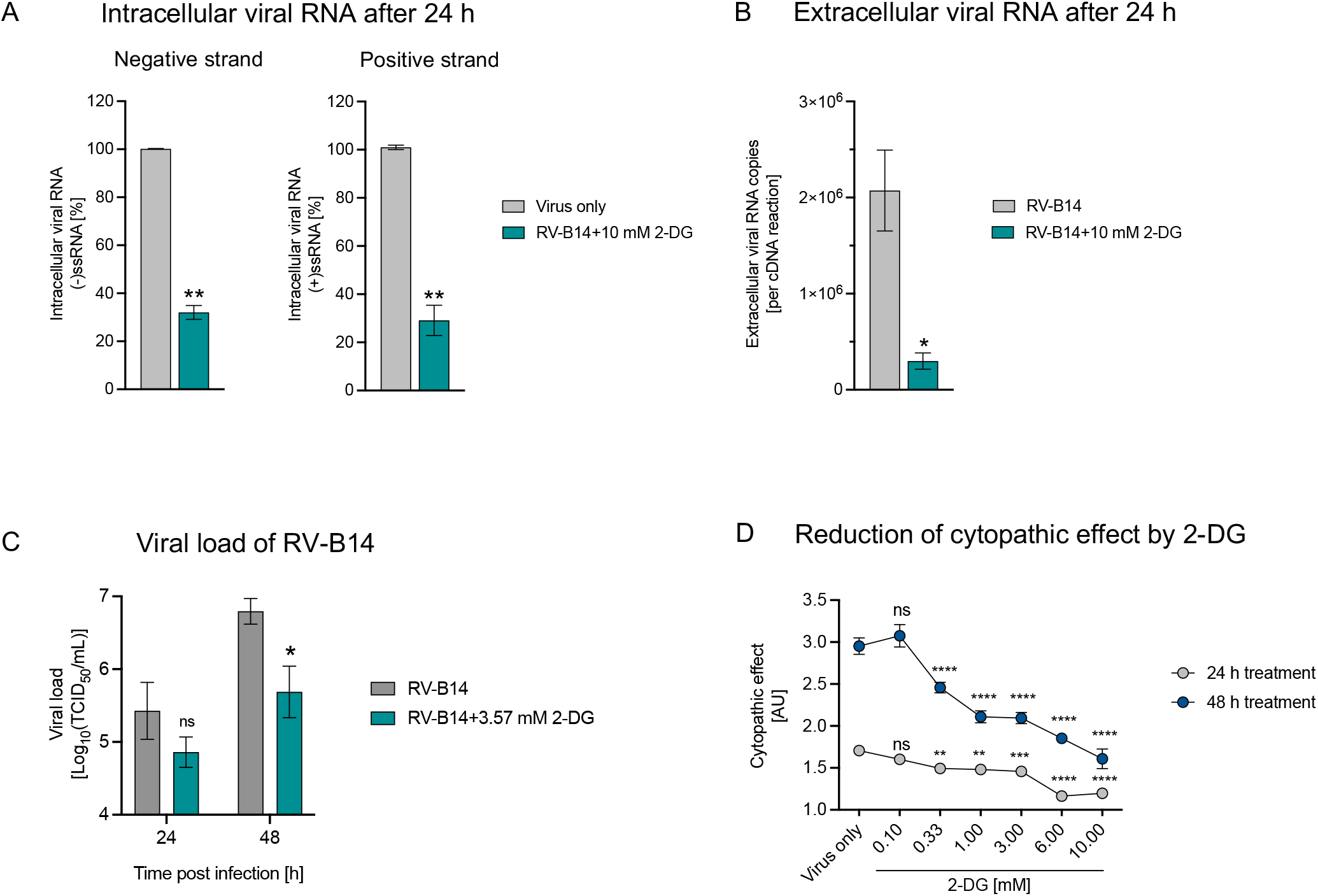
2-DG disrupts RNA template strand synthesis and inhibits RV-mediated cell death. HeLa Ohio cells were infected with RV-B14 (0.5 TCID_50_/cell) and treated with 10 mM 2-DG for 24 h to measure intracellular negative and positive viral RNA strand (A) or released extracellular viral RNA (B). Cells infected with RV-B14 (0.005 TCID_50_/cell) were treated with 3.57 mM 2-DG (IC_90_ for RV-B14) for up to 48 h at 34°C to measure viral load (C). Cells infected with RV-B14 (0.5 TCID_50_/cell) and treated with the indicated concentrations of 2-DG for 24 h or 48 h at 37 °C for measurement of virus-induced cytopathic effect (D). Graphs show pooled results ± SEM of 2-4 independent experiments (A,B,D) or one experiment (C). ns: non-significant; p < 0.05 (*p ≤ 0.05, **p ≤ 0.01, ***p ≤ 0.001, ****p ≤ 0.0001). RV: rhinovirus. AU: Arbitrary units

A characteristic of RV infection of tissue culture cells is the cytopathic effect (CPE) [39]. The impact of increasing concentrations of 2-DG on RV-induced cell death was assessed in HeLa Ohio cells at 24 h and 48 h post-infection. A significant reduction in CPE was seen in cells treated with 2-DG at 0.33 mM and higher after 24 h (Figure 3D). At 48 h post-infection, a stronger CPE could be observed in infected but untreated cells (‘Virus only’) and cell death was significantly reduced upon treatment with 2-DG at 0.33 mM or higher (Figure 3D). Together, these results suggest that 2-DG affects the RV life cycle by suppressing viral RNA replication and viral load and reduces RV-mediated cell death.

### 2-DG decreases CoV viral load

Similar to RVs, SARS-CoV-2 was recently shown to exploit the host glucose metabolism for replication and can potentially be targeted by 2-DG [24], [35]. However, 2-DG’s effect on endemic HCoVs hasn’t been investigated so far. With this rationale we investigated the effect of 2-DG on the viral load of the pandemic strain, SARS-CoV-2 as well as the two endemic human CoV stains, HCoV-229E and HCoV-NL63. Cells with known susceptibility to these coronaviruses were treated with increasing concentrations of 2-DG for 24 h to 48 h. The supernatant containing released virus was sampled every 24 h and viral load was assessed as TCID_50_. We observed a significant reduction in SARS-CoV-2 at 24 h post-infection at the highest tested 2-DG concentration (10 mM), and further, lower 2-DG concentrations led to significant effects 48h post-infection (Figure 4A). A similar behavior was observed for HCoV-229E, where 24 h and 48 h post-infection, a significant reduction in viral load was observed in cells treated with 0.32 mM and 1 mM 2-DG (Figure 4B). The use of lower 2-DG concentrations was based on decreased viability of MRC5 cells at 2-DG concentrations above 1 mM (data not shown). In the case of HCoV-NL63, there was no significant decrease in viral load at 24 h, however, at 48 h post-infection 2-DG concentrations above 1 mM suppressed viral load significantly (Figure 4C). These results suggest that 2-DG exerts a dose-dependent reduction in viral load of pandemic as well as endemic CoV strains.

**Figure 4:**
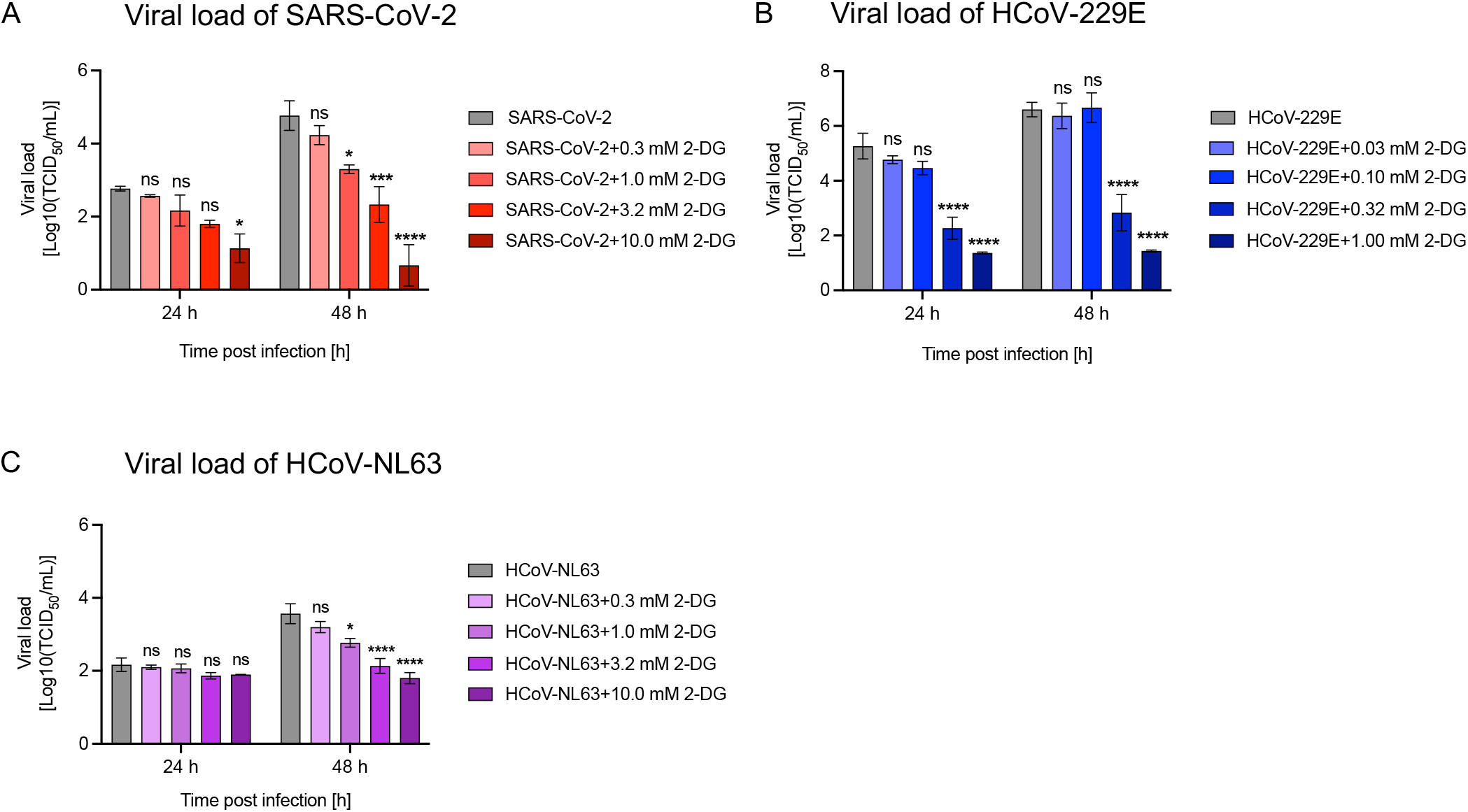
2-DG shows a dose-dependent antiviral effect on human coronaviruses. Viral load was measured from cell culture supernatants 24 h to 48 h post-infection. 2-DG treatment with the indicated concentrations was started 1 h post-infection. Viral load of SARS-CoV-2 (MOI 0.001) released from LLC-MK2 cells (A), HCoV-229E (MOI 0.01) released from MRC5 cells (B) and HCoV-NL-63 (MOI 0.01) released from LLC-MK2 cells (C). Graphs show pooled results ± SEM of 3 independent experiments. ns: non-significant; p < 0.05 (*p ≤ 0.05, **p ≤ 0.01, ***p ≤ 0.001, ****p ≤ 0.0001). SARS-CoV-2: severe acute respiratory syndrome coronavirus 2, HCoV: human coronavirus.

## DISCUSSION

In this study we investigated a host-directed approach to combat rhinovirus (RV) and coronavirus (CoV) infection by using 2-Deoxy-D-glucose (2-DG). This approach is based on the understanding that virus-induced metabolic reprogramming of the host cell plays a crucial role in viral replication [21], [22], [25]. Previously, Gualdoni et al. [25] demonstrated that 2-DG reverts RV-induced metabolic reprogramming of host cells and inhibits RV-B14 replication. Consequently, in the present study, we investigated the antiviral activity of 2-DG against additional minor- and major-group RVs, where 2-DG showed a dose-dependent inhibition of RV replication in epithelial cells including primary human nasal epithelial cells (HNECs). Simultaneously, we showed that treatment with 2-DG does not induce cytotoxic effects in this setting. Further, we sought to elucidate the implications of 2-DG on the RV replication cycle, intracellular kinetics of 2-DG and its impact on RV viral load. We found that 2-DG treatment led to a marked inhibition of template negative strand as well as genomic positive strand RNA replication. 2-DG treatment caused a significant reduction in the extracellular viral RNA level, RV viral load and in the RV-mediated cytopathic effect. At a physiological glucose concentration, 2-DG treatment led to enhanced inhibition of RV replication as compared to conventional high-glucose culture conditions. Assessment of 2-DG’s intracellular kinetics showed accumulation of the active intermediate, 2-DG6P, for several hours. Our concurrent study of 2-DG’s impact on CoVs also showed a significant reduction in viral load. Taken together, the results suggest 2-DG to be a potential broad-spectrum antiviral.

In our study, treatment with 2-DG inhibited replication of all tested minor- and major-receptor group strains of RV in HeLa Ohio cells under conventional culture condition (i.e., 2 g/L glucose) (Supplement Figure 1) and in primary human nasal epithelial cells (HNECs) (Figure 1C). As 2-DG competes with glucose for cellular uptake [26], [27], we lowered the glucose concentration to 1 g/L glucose – mimicking the human plasma glucose concentration – to assess the efficacy of 2-DG in a physiological context. We found that lower glucose concentrations potentiated 2-DG-mediated inhibition of RV replication, pointing to a higher efficacy of 2-DG in physiological settings (Figure 1A, Supplement table 2). It should be noted that the glucose concentration in fluid lining the nose and lung epithelium in humans is around 12.5 times lower than in plasma [40]. Therefore, it can be anticipated that 2-DG exhibits even higher antiviral efficacy in therapeutic target tissues. However, additional studies in models closer to the physiologic conditions are warranted to test this hypothesis. Further, as exposure to 2-DG has been shown to induce cytotoxic effects [41]–[43] we specifically tested the effect of 2-DG on HNECs and found no significant reduction in cell viability after 7 h 2-DG treatment (Figure 1D). Based on the experimental evidence and toxicology studies, the safety and pharmacokinetics of local (intranasal) 2-DG administration is currently being investigated in a Phase I clinical trial in Austria (NCT05314933) [44].

In the next step, we characterized the intracellular kinetics of 2-DG6P after a short exposure to 2-DG (Figure 2A). In the cell, 2-DG is phosphorylated to 2-DG6P, leading to its intracellular accumulation. Cytochalasin B, an inhibitor of the glucose transporter, was used as a control to ensure 2-DG6P specificity in our set-up (data not shown). Overall, we found that 2-DG6P was detectable up to several hours in HeLa Ohio cells and HNEC after a short incubation of the cells with 2-DG. The setup in this experiment mimics the *in vivo* setting where local treatment, e.g., in the nose, would only lead to a short exposure of epithelial cells to 2-DG. Our results suggest that even a brief exposure time is sufficient for extended inhibition of glycolysis via 2-DG6P and thereby to exhibit an antiviral effect.

During the RV replication cycle, the viral polyprotein is first generated via translation from the (+)ssRNA genome, which is then processed by viral proteases to generate viral proteins including the viral RNA polymerase [45]. Next, RNA polymerase generates (−)ssRNA strand copies, which in turn serve as a template for the multifold replication of the positive stand viral genome to be packaged in viral capsids, finally leading to release of the mature virions [46]. As conventional qPCR holds limitations to detect the negative strand in excess of positive strand copies, we employed a recently published strategy by Wiehler and Proud [32] to analyze the negative strand level. We observed that 2-DG significantly reduced the template (−)ssRNA as well as the genomic (+)ssRNA, a likely cause for the measured significant reduction in detectable extracellular viral RNA (Figure 4A&B). These findings point at a 2-DG-mediated impairment in viral RNA replication, resulting in a reduced amount of released virus. In line with this, TCID_50_ titration of the released virus on HeLa Ohio cells showed a reduction in viral load (Figure 4C). To be noted, HeLa Ohio cells used in this experimental setup, due to their cancerous origin, have a high glucose demand and are especially sensitive to glucose starvation and 2-DG treatment. Therefore, low amounts of 2-DG were used, and the cells were treated only once after the start of the RV infection. This could explain the relatively small difference in viral load (Figure 3C) in contrast to the significant difference in released extracellular viral RNA (Figure 3B).

In our subsequent analysis, we found that 2-DG exerted a protective effect by significantly reducing virus-induced cell death in HeLa Ohio cells (Figure 4D). In contrast, RV infection does not cause cell lysis in cultures of healthy bronchial epithelial cells [47]. Interestingly, the same study reported increased viral replication and cell lysis after RV infection in asthmatic bronchial epithelial cells [47]. Based on these findings, we could envision protection of RV-infected bronchial epithelial cells of asthma patients by 2-DG.

The host metabolic dependency of CoVs is similar to that of RVs and studies suggest that 2-DG alters SARS-CoV-2 replication [24], [26], [48]. These results prompted us to further investigate the effect of 2-DG on CoVs infection. In our study, 2-DG treatment of pandemic SARS-CoV-2 resulted in a dose-dependent reduction of viral load. In line, 2-DG has been approved for use in patients with moderate to severe SARS-CoV-2 infection in India by Drug Controller General India (DCGI) after performance of Phase II and Phase III clinical trials conducted by the Defense Research and Development Organization (DRDO), India in collaboration with Dr Reddy’s Laboratories, India [49]. However, the peer reviewed data of the trials are still unpublished. Further, in our study, we show for the first time the antiviral effect of 2-DG on endemic HCoVs 229E and NL63. As in the case of SARS-CoV-2, 2-DG caused a dose-dependent reduction in viral load in both endemic HCoV strains.

Comparing our data from RV viral load, lower concentrations of 2-DG are sufficient to cause a long-term significant reduction in viral load in both endemic and pandemic CoVs. The difference between RV and CoV with respect to the required 2-DG concentrations can be attributed to differences in cell culture models. Another possible explanation is that CoVs are enveloped [13] and contain glycosylated envelope proteins responsible for host cell interaction and infection. Along with CoVs dependence on host glucose metabolism for replication [24], they are also dependent on the host cell machinery for glycosylation of viral proteins [50]. Thus, the reduction in CoV viral load could originate from 2-DG not only inhibiting glycolysis but also affecting protein and lipid glycosylation [51]. However, further studies are required to decipher a possible role of 2-DG in the production of defective virions in enveloped viruses.

In conclusion, we present further *in vitro* data that support a host-directed approach to tackle RV and CoV infections. The dependency of these viruses on the host cell metabolism and cell machinery reveals a therapeutic opportunity to target them with host-directed antivirals such as 2-DG. The low cytotoxicity of 2-DG and the long half-life of the active metabolite 2-DG6P advocates its short-time topic application at comparably high concentrations, e.g., as a spray to be employed early in infection, which might safely block viral spreading.

## ACKNOWLEDGMENTS

We thank Melanie Graf and the Global Pathogen Safety Team (Takeda), most notably Jasmin de Silva, Elisabeth List and Effie Oindo (experiments), Veronika Sulzer (cell culture), Eva Ha, Simone Knotzer and Alexandra Schlapschy-Danzinger (virus propagation). SARS-CoV-2 was sourced via EVAg (supported by the European Community) and kindly provided by Christian Drosten and Victor Corman (Charité Universitätsmedizin, Institute of Virology, Berlin, Germany). HCoV-NL63 was kindly provided by Lia van der Hoek (Medical Microbiology, Academisch Medisch Centrum, Amsterdam, Netherlands).

## AUTHOR CONTRIBUTION

L.W., M.K., S.C., V.K., A.A., X.C., D.S. and A.-D.G. performed experiments and analyzed data. D.B. and I.G. provided virus strains, reagents, and valuable input. A.-D.G., J.S., T.R.K., M.K. and G.G. were in charge of planning and directing the study. L.W and A.-D.G. wrote the manuscript with input from co-authors. All authors read and approved the final manuscript.

## SUPPLEMENTARY MATERIALS

**Supplement figure 1: Inhibition of RV replication by 2-DG in HeLa Ohio cells.** Cells were cultivated in medium containing 2 g/L glucose. Intracellular viral RNA was assessed by qPCR 7 h post-infection at 0.005 TCID_50_/cell for the indicated RV strains in HeLa Ohio cells. Cells were treated with the indicated concentrations of 2-DG (represented on a log10 scale) 1 h post-infection until the samples were collected. Graphs show pooled results ± SEM of 3 independent experiments. RV: rhinovirus.

**Supplement table 1:** Materials used in the study.

**Supplement table 2:** IC_50_ values of tested RV strains in HeLa Ohio and HNECs.

