## Supplemental Table 1 for "Host-directed therapy with 2-Deoxy-D-glucose inhibits human rhinoviruses, endemic coronaviruses, and SARS-CoV-2"

**Supplementary Table1:** Materials

**Cell lines, cell culture medium and virus stocks**

| **Item** | **Company** | **Cat.No.** | **Additional Information** |
| --- | --- | --- | --- |
| MRC-5 cells | ATCC | CCL-171 |  |
| LLC-MK2 cells | ATCC | CCL-7 | Amsterdam UMC |
| Vero cells | ATCC | CCL-81 | sourced from ECACC (84,113,001) |
| Human nasal epithelial cells (HNECs) | Epithelix Sàrl | EP51AB | Donor no. AB0630.01 |
| HeLa Ohio cells | ECACC | 14J003 |  |
| MRC-5 cell culture media | Gibco – Life Technologies | 31095 | Eagle-MEM supplemented with 10 % FBS, L-glutamine (2 mM), non-1x essential amino acids solution, 1 mM sodium pyruvate, 100 mg/mL Gentamycin sulfate and 0.15 % sodium bicarbonate. |
| LLC-MK2 cell culture media | Gibco – Life Technologies | 31095 | Eagle-MEM supplemented with 10 % FBS, 1x non-essential amino acids solution, 100 mg/mL Gentamycin sulfate and 25 mM HEPES. |
| Vero cell culture media | Takeda Inhouse |  | TC-Vero medium supplemented with 5 % FBS, 2 mM L-glutamine, 1x non-essential amino acids solution, 1 mM sodium pyruvate, 100 mg/mL Gentamycin sulfate and 0.075 % sodium bicarbonate. |
| Human nasal epithelial cell culture medium | STEMCELL Technologies | 05041, 07925, 15140122 | PneumaCult™-Ex Plus Basal Medium supplemented with 1x PneumaCult™-Ex Plus Supplement, 0.1 % Hydrocortisone Stock Solution and 1 % Penicillin/Streptomycin (100 Units/ml). |
| HeLa Ohio cell culture medium | Gibco | 21875034, 11879020 | RPMI 1640 medium supplemented with 10 % FBS, 1 % Penicillin/Streptomycin (100 Units/ml) and 2 mM L-glutamine. |
| Glucose-free medium | Gibco | 11879020 | RPMI 1640 medium without glucose containing 10 % FBS, 1 % Penicillin/Streptomycin (100 Units/ml) and 2 mM L-glutamine. |
| TCID_50_ medium | Gibco |  | MEM-EARLES supplemented with 2 % FBS, 1 % Penicillin/Streptomycin (100 Units/ml), 2 mM L-glutamine and 30 mM MgCl_2_. |
| HCoV-229E | ATCC | VR-740 | Rockville, MD |
| HCoV-NL63 | Lia van der Hoeck/ Laboratory of Experimental Virology Dep Medical Microbiology |  | Amsterdam UMC  Das AMC wird vor Veröffentlichung der Daten informiert und ggf. wird Autorenschaft angeboten. |
| SARS-CoV-2 strain BetaCoV/Germany/BavPat1/2020 | EVAg | 026V-03883 | provided by the Charité Universitätsmedizin, Institute of Virology, Berlin, Germany |
| HRV-A54 | ATCC | VR1661 | Prepared by Irene Gösler, Blaas group (MPL, Vienna, Austria) |
| HRV-B14 | ATCC | VR284 | Prepared by Irene Gösler, Blaas group (MPL, Vienna, Austria) |
| HRV-A1B | ATCC | VR1645 | Prepared by Irene Gösler, Blaas group (MPL, Vienna, Austria) |
| HRV-A2 | ATCC | VR482 | Prepared by Irene Gösler, Blaas group (MPL, Vienna, Austria) |
| HRV-A89 | ATCC | VR1199 | Prepared by Irene Gösler, Blaas group (MPL, Vienna, Austria) |
| HRV-A16 | ATCC | VR283 | Prepared by Irene Gösler, Blaas group (MPL, Vienna, Austria) |

**Chemicals, equipments, reagents and softwares**

| **Item** | **Company** | **Cat.No.** |
| --- | --- | --- |
| PBS | Takeda in-house | 7500144 |
| D-PBS | Gibco | 17-516F |
| MycoZAP | Lonza | VZA-2032 |
| FCS | Gibco/Corning | 10270106/35070CV |
| L-glutamine | Gibco/Thermo Fisher | 25030024 |
| Penicillin- Streptomycin | Gibco | 15140122 |
| Gentamicin | Gibco/Thermo Fisher | 15710049 |
| Sodium pyruvate | Gibco/Thermo Fisher | 11360039 |
| Non-essential amino acid solution (NEAS) | Gibco/Thermo Fisher | 11140035 |
| Sodium bicarbonate | Gibco/Thermo Fisher | 25080060 |
| HEPES | Gibco/Thermo Fisher | 15630056 |
| Animal Component Free cell dissociation kit | STEMCELL Technologies | 5426 |
| Accutase | Corning | 25058049 |
| 2-DG | Sigma | D8375-5G |
| Triton X-100 | Roche | 33981126 |
| Crystal violet powder | Acros Organics | 405830250 |
| Methanol (>= 99.9%) | Fisher Chemical |  |
| Acetic acid glacial (>= 99.7%) | Fisher Chemical | A/0400/PB08 |
| T25 cell culture flasks | Corning | 430639 |
| Nunc™ EasYFlask™ cell  culture flask T75 | Thermo Scientific | 10364131 |
| 96well plates | Corning | 3598 |
| Deep-well plates | Greiner | 780271 |
| Costar® TC-  treated 24 well plate | Corning | CLS3527 |
| MicroAmp 96-well microtiter plate | Applied Biosystems | I02K0 Q415 |
| Adhesive seals for PCR microtiter plates | Thermo Scientific | AB1170 |
| Nuclease free water | Neofroxx | 1058LT001 |
| First strand cDNA  Synthesis | Fischer Scientific | K1612 |
| PowerTrackTM SYBR Green Master Mix | Applied Biosystems | A46109 |
| ExtractMe Total RNA Kit+DNAse I | Blirt | EM31 |
| Universal mouse reference RNA (spike-in control) | Invitrogen | QSO640 (14920) |
| Glucose-Uptake Glo^TM^ Assay kit | Promega | J1341 |
| Microscope | Nikon | Diaphot 200 or Eclipse TS100 |
| CASY® Cell Counter and Analyzer TT | Biovendis | N.a. |
| QuantStudio 1Real-Time PCR Instrument | Applied Biosystems | A40425 |
| Nanophotometer N50-Touch | IMPLEN | T51384 |
| GENios, GENios FL and GENios Plus microplate reader | Tecan |  |
| GraphPad Prism |  |  |

**Primers and oligo sequences**

| **Name** | **Sequence (5’-3’)** |
| --- | --- |
| HPRT_F | TCAGGCAGTATAATCCAAAGATG |
| HPRT_R | AGTCTGGCTTATATCCAACACTT |
| HRV-A54_F | CCTGTGAGCCCTCTGAAAAG |
| HRV-A54_R | GGGGATTGCAAGTCATCTGT |
| HRV-B14_F | GGCGCCATATCCAATGGTGT |
| HRV-B14_R | TCCACCTGATCGAACGTCCA |
| HRV-A1B_F | TCTACGCGCAGCAGATAATG |
| HRV-A1B_R | ACTGCCACTGGCTCACTTCT |
| HRV-A2_F | TAATGTGGCAGCAGCGTTTC |
| HRV-A2_R | ATGTTGTACACCTGCGCAAG |
| HRV-A89_F | GCAATGCTAAGTGCTGTCCA |
| HRV-A89_R | AGGTGGAGGAGATTGGAGGT |
| HRV-A16_F | CCAAACACACCCAATACTGCTG |
| HRV-A16_R | TTGGTCCAGTTTGCTTGTGC |
| HRV-B14 primer amplicon | GGCGCCATATCCAATGGTGTCTATGTACAAGCACTTCTGTTTCCCAGGAGCGAGGTATAGGCTGTACCCACTGCCAAAAGCCTTTAACCGTTATCCGCCAACCAACTACGTAACAGTTAGTACCATCTTGTTCTTGACTGGACGTTCGATCAGGTGGA |
| mZfp866_F2 | ATC CAG GTG AGC TGA GGG AT |
| mZfp866_R2 | AAC AGC TGA GCC ATC TCA CC |
| chimHRV-b14_RT | ATCAGCGATGCCGAACGTATGGCGCCATATCCAATGGTGT |
| chimHRV-b14_R1 | ATCAGCGATGCCGAACGTAT |
