## Supplemental Table 2 for "Host-directed therapy with 2-Deoxy-D-glucose inhibits human rhinoviruses, endemic coronaviruses, and SARS-CoV-2"

**Supplementary Table 2:** Summary of absolute half-maximal inhibitory concentration (IC_50_)

| **Cell type** | **Virus strain** | **Glucose content [g/L]** | **Absolute IC50 [mM]** |
| --- | --- | --- | --- |
| HeLa Ohio cells | RV-B14 | 2 | 3.44 |
|  |  | 1 | 1.15 |
|  | RV-A89 | 2 | 9.08 |
|  |  | 1 | 2.36 |
|  | RV-A16 | 2 | 9.22 |
|  |  | 1 | 2.67 |
|  | RV-A54 | 2 | 7.97 |
|  |  | 1 | 2.08 |
|  | RV-A2 | 2 | 6.08 |
|  |  | 1 | 2.18 |
|  | RV-A1B | 2 | 6.87 |
|  |  | 1 | 1.98 |
| Undifferentiated HNECs | RV-B14 | Not disclosed | 7.77 |
|  | RV-A16 |  | 6.35 |
