## Supplementary figures and images for "Host-directed therapy with 2-Deoxy-D-glucose inhibits human rhinoviruses, endemic coronaviruses, and SARS-CoV-2"

### Supplemental Figure 1

Supplement Figure 1

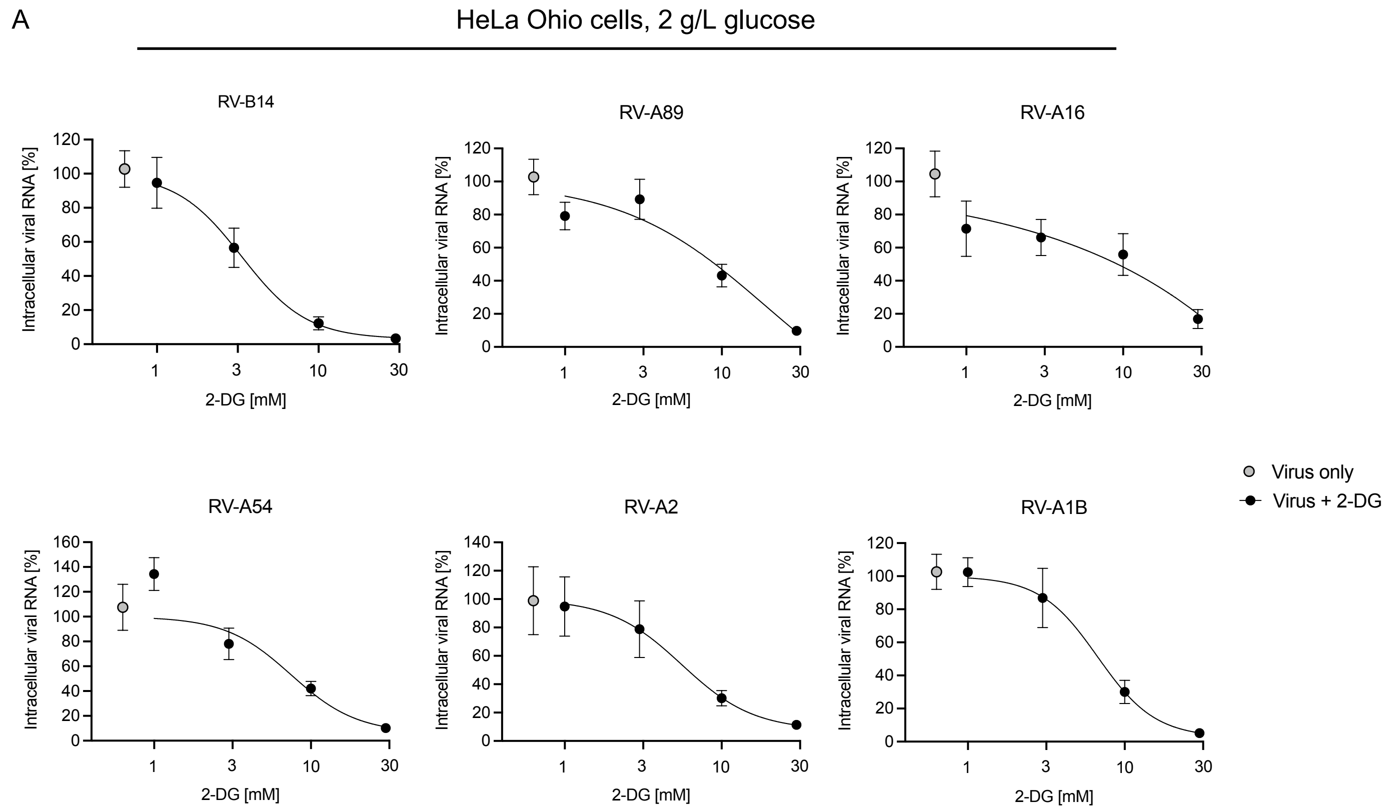
